# Recruitment of an ancient branching program to suppress carpel development in maize flowers

**DOI:** 10.1101/2021.09.03.458935

**Authors:** Harry Klein, Joseph Gallagher, Edgar Demesa-Arevalo, María Jazmín Abraham-Juárez, Michelle Heeney, Regina Feil, John E. Lunn, Yuguo Xiao, George Chuck, Clinton Whipple, David Jackson, Madelaine Bartlett

## Abstract

Floral morphology is immensely diverse. One developmental process acting to shape this diversity is growth suppression. For example, grass flowers exhibit extreme diversity in floral sexuality, arising through differential suppression of stamens or carpels. In maize, carpels undergo programmed cell death in half of the flowers initiated in ears and in all flowers in tassels. The HD-ZIP I transcription factor gene *GRASSY TILLERS1* (*GT1*) is one of only a few genes known to regulate this process. To identify additional regulators of carpel suppression, we performed a *gt1* enhancer screen, and found a genetic interaction between *gt1* and *ramosa3* (*ra3). RA3* is a classic inflorescence meristem determinacy gene that encodes a trehalose-6-phosphate (T6P) phosphatase (TPP). Dissection of floral development revealed that *ra3* single mutants have partially derepressed carpels, whereas *gt1; ra3* double mutants have completely derepressed carpels. Surprisingly, *gt1* suppresses *ra3* inflorescence branching, revealing a role for *gt1* in meristem determinacy. Supporting these genetic interactions, GT1 and RA3 proteins colocalize to carpel nuclei in developing flowers. Global expression profiling revealed common genes misregulated in single and double mutant flowers, as well as in derepressed *gt1* axillary meristems. Indeed, we found that *ra3* enhances *gt1* vegetative branching, similar to the roles for the trehalose pathway and *GT1* homologs in the eudicots. This functional conservation over ~160 million years of evolution reveals ancient roles for *GT1-*like genes and the trehalose pathway in regulating axillary meristem suppression, later recruited to mediate carpel suppression. Our findings expose hidden pleiotropy of classic maize genes, and show how an ancient developmental program was redeployed to sculpt floral form.

## Introduction

Variation in development drives variation in organismal form. One important process in floral development and evolution is growth suppression in floral organs (1, 2). A prominent form of this suppression exists in the grass family (Poaceae). Most grass flowers initiate both carpel and stamen primordia, but selective suppression of these primordia has led to immense diversity in floral sexuality (3). This diversity is critical for patterns of gene flow in natural populations, fertile flower production in nature and agriculture, and facilitates controlled crosses in breeding programs (4–6). Despite these important consequences to both evolution and agriculture, only a handful of genes are known to regulate floral sexuality in the grasses.

Floral sexuality has long been studied in *Zea mays* (maize), where programmed cell death suppresses carpel development in all tassel flowers, and in one of the two flowers in each ear spikelet (7, 8). Among the few characterized carpel suppression genes, most encode enzymes with roles in hormone metabolism and have pleiotropic effects on development and defense (9–15). This list includes several genes underlying the classic *tasselseed* (*ts*) mutants, which exhibit extensive tassel feminization beyond carpel suppression (9, 10, 12–14, 16). Most of the cloned *ts* mutants encode genes involved in jasmonic acid metabolism, whereas *ts4* encodes a miRNA which targets *Ts6*, a developmental regulator with multiple roles in flower and inflorescence development (12, 14).

Most genes that affect carpel suppression simultaneously affect other traits that differentiate tassels from ears, such as stamen development, bract (glume) morphology, and inflorescence morphology (9–11, 16). *GRASSY TILLERS1* (*GT1*) is an exception. While *GT1* was defined by its role in regulating axillary branching, *gt1* mutants have a weak carpel suppression phenotype in otherwise normal flowers and inflorescences (17). To find additional regulators of carpel suppression, we conducted an enhancer screen of *gt1* mutants and found a genetic interaction with the classic inflorescence determinacy gene, *RAMOSA3* (*RA3*) (18). Our results reveal surprising pleiotropy and interactions between *gt1* and *ra3*, which together regulate carpel suppression, meristem determinacy, and axillary meristem suppression.

## Results

### The *rapunzel* (*rzl*) genes suppress carpels in tassel and ear flowers

The mutants we identified in our *gt1* enhancer screen had long silks emerging from tassels, so we called this phenotype *rapunzel* (*rzl*), after the Grimm brothers’ fairy tale character with long hair. Two of these double mutants, *gt1;rzl-3* and *gt1;rzl-4,* phenocopied one-another, and did not complement, indicating that *rzl-3* and *rzl-4* were allelic (Table S1).

The *gt1;rzl-3* and *gt1;rzl-4* floral phenotypes were specific to carpels. In contrast to most *tasselseed* mutants (9, 10, 12, 13), stamen development in *gt1;rzl-3* and *gt1;rzl-4* tassel flowers was not suppressed (Fig. 1, Table S2). Except for a difference in carpel suppression in tassel flowers, *rzl-3* mutants (*rzl-3; gt1/+*) were indistinguishable from *gt1* single mutants (Fig. 1H-I). In normal ear spikelets, two flowers initiate, but only the upper flower completes development; the lower flower is suppressed (7). In *gt1;rzl-3* and *gt1;rzl-4* ears, carpel growth was derepressed in lower flowers, resulting in two fertile flowers per spikelet (Fig. 1F, Fig. S1). Thus, *rzl-3/4* interacts with *gt1* to disrupt carpel suppression, while leaving other floral and inflorescence traits unaffected.

**Figure 1.**
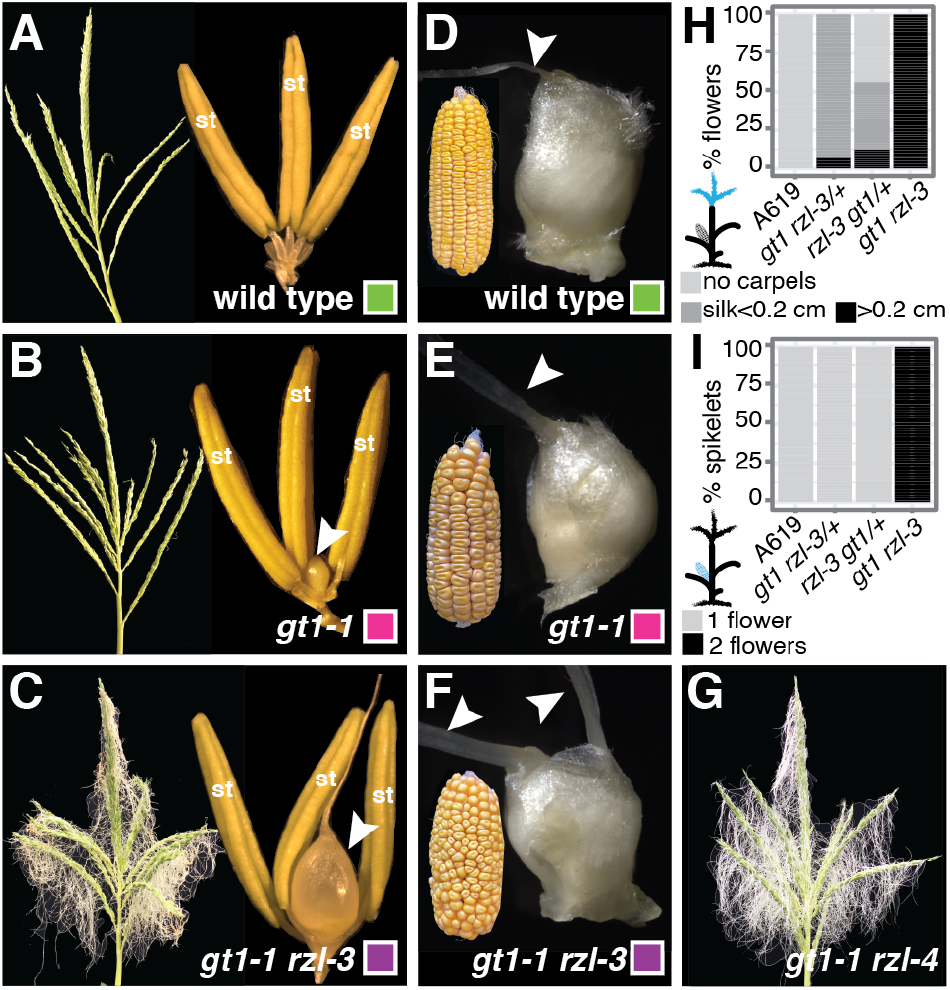
*gt1; rzl* double mutants have derepressed carpels in tassels and ears. (**A-C**) tassels and tassel flowers, and (**D-F**) ears and ear flowers in (**A,D**) wild type (A619), (**B,E**) *gt1; rzl-3/+*, (**C,F**) *gt1; rzl-3*, and (**G**) *gt1; rzl-4* mutants. Arrowheads indicate derepressed gynoecia. (**H-I**) Gynoecia are derepressed in *gt1; rzl-3* tassel and ear flowers.

### *rzl-3* and *rzl-4* are alleles of the trehalose-6-phosphate phosphatase gene, *RAMOSA3*

To identify the *rzl-3/4* gene, we used both bulked segregant analysis coupled with whole genome shotgun Categorization of (**H**) tassel flowers (20 flowers from each of 5 individuals per genotype) and (**I**) ear spikelets (50 spikelets from each of 5 individuals per genotype). st = stamen sequencing (BSA-seq), and fine mapping (19). Our BSA-Seq results revealed a broad peak between 160 and 180 Mbp on chromosome seven that represented the *rzl-4* mapping interval (Fig. 2). The narrow peaks on chromosomes three and nine are likely because of differences between B73 lab stocks and the reference genome (19, 20). The broad region of homozygosity on chromosome 1 results from the introgression of *gt1*, which arose in A619 (17), into B73. Using fine mapping (19), we reduced the chromosome 7 mapping interval to a ~230 kbp region containing seven genes, only one of which contained a canonical EMS SNP predicted to negatively affect gene function (21).

**Figure 2.**
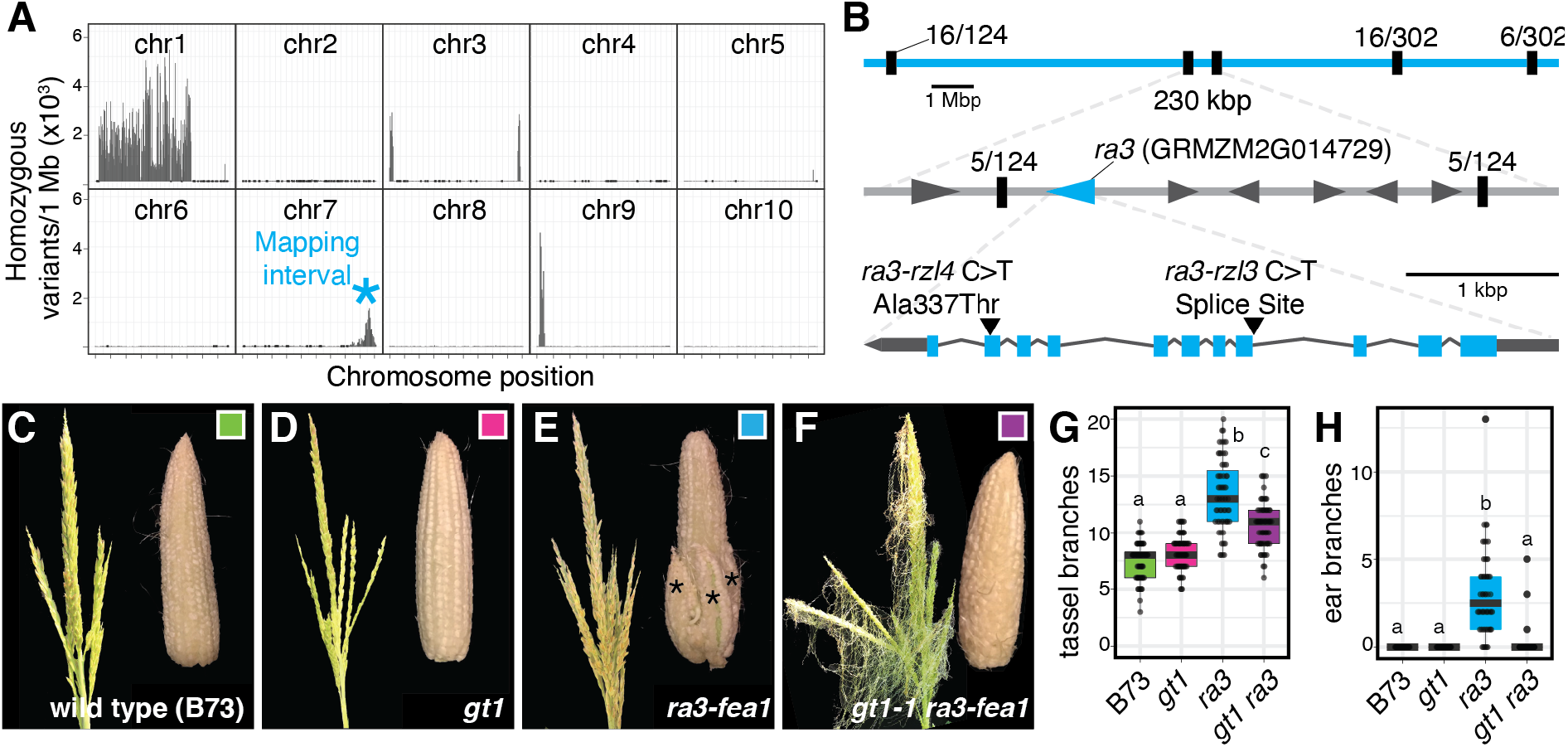
*rzl-3* and *rzl-4* are alleles of *ramosa3* (*ra3*), which interacts genetically with *gt1* in flowers and inflorescences. (**A**) BSA-Seq mapped *rzl-4* to chromosome 7. (**B**) Mapping interval shown in detail. Black tick marks indicate genetic markers, numbers above ticks indicate recombinants identified in fine mapping. (**C-F**) Tassels and ears from (**C**) B73 (wild type), (**D**) *gt1*, (**E**) *ra3-fea1* mutants, and (**F**) *gt1; ra3-fea1* mutants, all in B73. (**G-H**) *gt1* suppresses *ra3* inflorescence branching in both (**G**) tassels and (**H**) ears. Different letters indicate statistically significant differences (p < 0.001, ANOVA, Tukey’s post-hoc).

This EMS SNP was predicted to change a single amino acid (Ala337Thr) in the trehalose-6-phosphate phosphatase (TPP) encoded by *RAMOSA3 (RA3)* (18). Ala337 is deeply conserved in TPP paralogs, and contacts the active site in homology models (Fig. S2). In addition, the substitution of a homologous amino acid in a RA3 paralog negatively affects protein function (22), suggesting that the *rzl-4* SNP would impact RA3 function similarly. Because *rzl-4* and *rzl-3* were allelic, we sequenced *RA3* in *rzl-3* mutants and found an EMS mutation at a likely splice acceptor site 5’ of exon 4. Transcripts from this *ra3* allele had an in-frame deletion of two active-site amino acid codons (Fig. S2). Thus, both *gt1;rzl-3* and *gt1;rzl-4* double mutants harbor alleles of *ra3* predicted to negatively affect gene function, indicating that *rzl-3* and *rzl-4* encode alleles of *ra3*. From now on we will refer to these alleles as *ra3-rzl3* and *ra3-rzl4.*

### *gt1* suppresses *ra3* tassel and ear branching

Usually, *ra3* mutants have branched ears and increased tassel branching due to indeterminate meristems that produce many spikelets on long branches (18, 22). Therefore, we were surprised to find unbranched ears in *gt1;ra3-rzl3/4* double mutants, and in *ra3-rzl3/4* single mutants heterozygous at *gt1* (Fig. 1). Given that *gt1* is semi-dominant, the lack of ear branching in these mutants could have been caused by a second genetic interaction between *gt1* and *ra3,* regulating meristem determinacy. The lack of ear branching may also have been the result of the genetic background (A619) used for EMS mutagenesis. Most characterization of *ra3* has been in the B73 genetic background (18, 22), and background modifiers affect the *ra3* ear determinacy phenotype (18).

To test for a genetic interaction between *gt1* and *ra3* independent of any A619 modifiers, we made a *gt1; ra3* double mutant with a third, well-characterized allele of *ra3* (*ra3-fea1*) in the B73 genetic background (18, 22). *gt1; ra3-fea1* double mutants in B73 recapitulated the *rzl* phenotype, with silks in tassel flowers, as in our *gt1; ra3-rzl3* and *gt1; ra3-rzl4* mutants (Fig. 2F). *ra3-fea1* single mutants had branched ears and increased tassel branching, as expected (18). However, most *gt1;ra3-fea1* double mutants lacked ear branches and had fewer tassel branches than *ra3* single mutants (Fig. 2C-H), indicating a second genetic interaction between *gt1* and *ra3,* regulating meristem determinacy. Thus, *RA3* and *GT1* act to regulate both meristem determinacy in inflorescences and carpel suppression in flowers.

### *ra3* mutants have a carpel suppression phenotype and RA3 colocalizes with GT1 in carpel nuclei

*ra3; gt1/+* mutants had a weak floral phenotype (Fig. 1), possibly because of *gt1* semi-dominance (17), or because *ra3* single mutants have a weak floral phenotype. Indeed, there have been hints of likely background-dependent *ra3* floral phenotypes (23). To further investigate the *ra3* floral phenotype, we followed tassel flower development in mutants using scanning electron microscopy.

In wild type flowers, the two silk carpels are first visible as a raised line of tissue called the gynoecial ridge. In ear flowers, these silk carpels grow to enclose the ovulate carpel, and fuse to form the stigma, called the silk in maize (7). In tassel flowers, carpels initiate, but undergo programmed cell death shortly after the initiation of the gynoecial ridge (8, 24) (Fig. 3A-D). In contrast, in *gt1* single mutants, silk carpels continued to grow past the gynoecial ridge stage and formed a peak of tissue over the developing ovulate carpel (Fig. 3E-H). The tassel flowers of *ra3* single mutants had carpels that resembled those of *gt1* mutants: the two silk carpels formed a peak of tissue over the ovulate carpel, but did not fuse laterally to enclose the ovulate carpel (Fig. 3I-L). These data indicate that carpel suppression is partially disrupted in both *ra3* and *gt1* single mutants.

**Figure 3.**
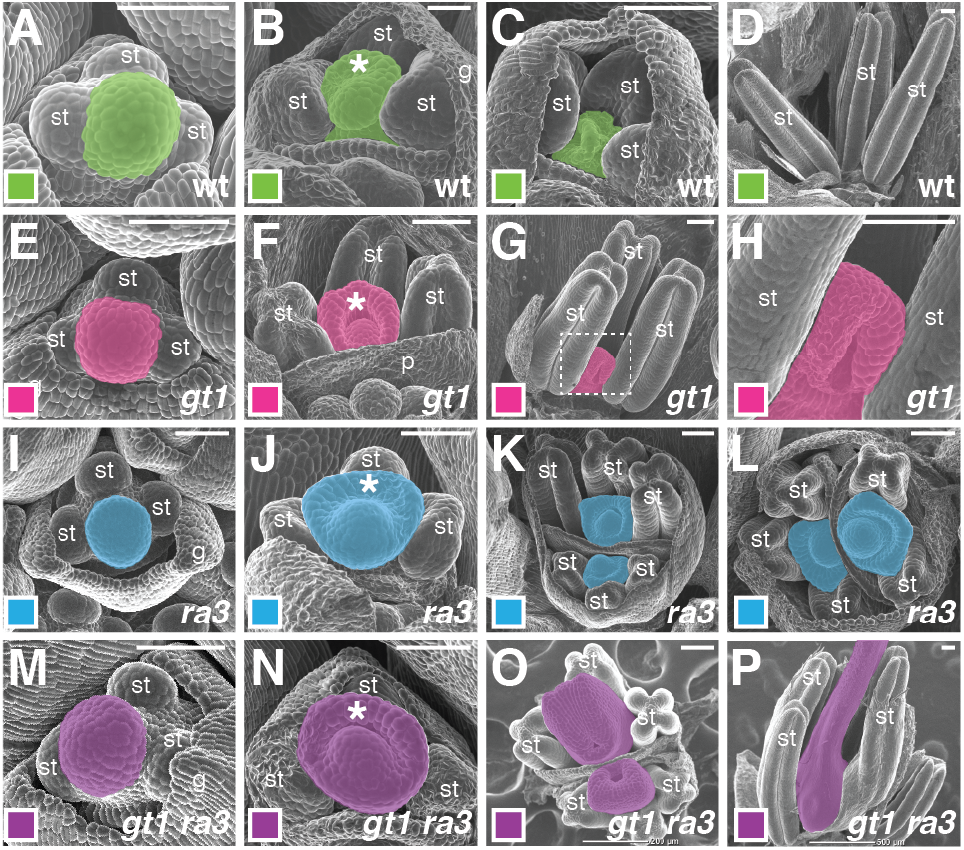
*ra3* single mutants exhibit a weak carpel derepression phenotype. Tassel flower development in (**A-D**) wild type (A619), (**E-H**) *gt1*, (**I-L**) *ra3*, and (**M-P**) *gt1; ra3* mutants. Outlined region in G shown at higher magnification in H. Gynoecia false-colored, no remains of gynoecium evident in D. g = glume, p = palea, st = stamen, * = gynoecial ridge.

The carpels in *gt1; ra3* double mutant flowers continued to grow well past the gynoecial ridge stage, eventually fusing to form silks (Fig. 3M-P, Fig. S1). In addition, the gynoecia of *gt1;ra3* double mutants closely resembled the gynoecia of wild type ear florets, with two silk carpels and an ovulate carpel (Fig. 3O, Fig. S1). In contrast, other floral meristem determinacy mutants, like *zea mays agamous1* (*zag1*), *bearded ear (bde), Tasselseed6/indeterminate spikelet1* (*Ts6/ids1*), *drooping leaf1* (*drl1*) and *drooping leaf2* (*drl2*), have indeterminate floral meristems, leading to disorganized gynoecia with extra primordia, many of which do not fuse to form silks (12, 25–29). Thus, our results indicate that the *gt1; ra3* phenotype arises not because the floral meristems had become indeterminate, but because floral organ suppression was disrupted.

The floral phenotypes of both single and double mutants led us to assess GT1 and RA3 localization in developing flowers, using immunofluorescence with custom antibodies. GT1 and RA3 both localized to the nuclei of carpel primordia in 1.5 cm tassels, prior to carpel growth suppression (Fig. 4A). Importantly, both GT1 and RA3 were predominantly localized to the nuclei of sub-epidermal cells, where programmed cell death initiates in maize and its relatives in the Andropogoneae (8, 30). While *RA3* likely acts non-cell autonomously in regulating meristem determinacy (18, 31), the GT1 and RA3 localization patterns in tassel flowers suggest that both act cell-autonomously in regulating carpel suppression.

**Figure 4.**
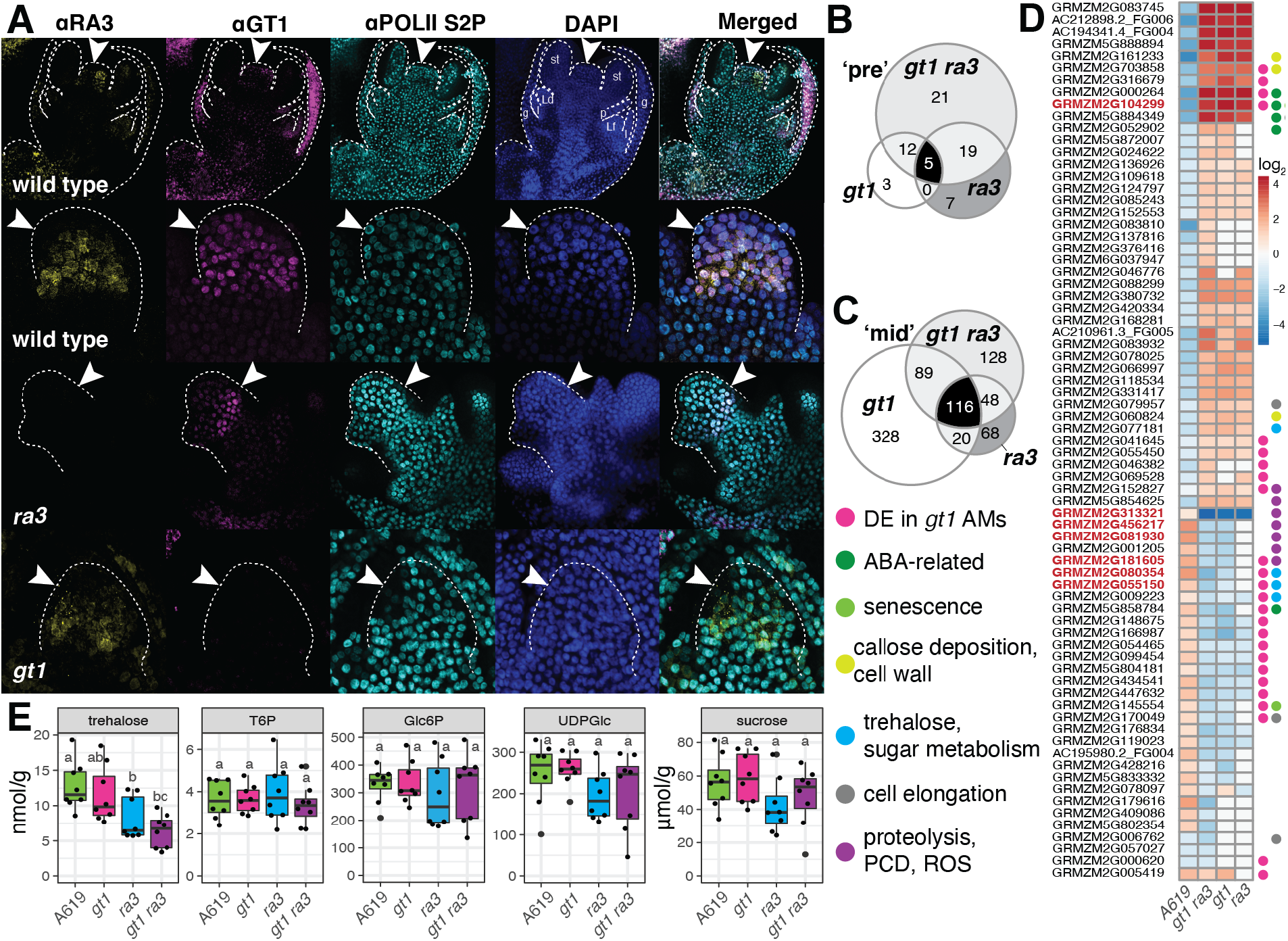
GT1 and RA3 proteins localized to nuclei of carpels in tassel flowers, where trehalose metabolism is impacted in *gt1; ra3* mutants. (**A**) fluorescent immunolocalizations of GT1, RA3, and aPOLII proteins using native antibodies in wild type, *gt1*, and *ra3* mutants. Secondary antibodies were labeled with Alexa 488 (aRA3), Alexa 647 (aGT1) or Alexa 568 (aPOL II). g = glume, l = lemma, Ld = lodicule, Lf = flower floret, p = palea, st = stamen. Arrowheads indicate gynoecia. Differentially expressed (DE) genes (logFC >= 1 and FDR < 0.05) in double and single mutant tassel primordia just prior to carpel suppressioin (**B**) and mid-suppression (**C**). (**D**) 73 genes misregulated in *gt1; ra3* tassels at the mid suppression stage also changed over time in wild type (A619) tassels. 26 of these genes are also misexpressed in *gt1* tiller buds (45). Genes discussed in text in red. (**E**) T6P and sucrose levels were not significantly different between A619 and mutant tassels. Trehalose was significantly lower in *gt1; ra3* tassels. Different letters indicate statistically significant differences (p < 0.05, ANOVA, Tukey’s post-hoc).

### GT1 and RA3 regulate genes predicted to mediate programmed cell death

To identify the effectors of carpel suppression downstream of *GT1* and *RA3*, we examined the transcriptional profiles of wild type (A619), *gt1, ra3,* and *gt1; ra3* tassels at two developmental timepoints: (1) prior to carpel suppression (pre-suppression, 0.8 - 1 cm tassels), and (2) during active carpel suppression (mid-suppression, 1 - 2 cm tassels). We reasoned that genes that were (1) misexpressed in *gt1; ra3* mutant flowers, and (2) changed over wild type carpel development represented the best carpel suppression gene candidates. We performed differential expression analyses on our sequencing data (32), and focused on the 73 genes that satisfied both of these conditions, and were highly differentially expressed at the mid-suppression stage (|fold change| > 2, Fig. 4B-D, Dataset S1).

In this set of carpel suppression candidate genes, most genes that were downregulated in mutants were upregulated over the course of development in A619, or vice-versa (Fig. 4D). Although the carpel suppression gene set was not enriched for specific GO-terms, it did contain several developmental regulators whose homologs in *Arabidopsis thaliana* (arabidopsis) have roles in reactive oxygen species (ROS) signaling, callose deposition and programmed cell death (Table 1). For example, homologs of the arabidopsis NAC transcription factor genes *KIRA1* (GRMZM2G081930) and *ANAC087* (GRMZM2G181605) were upregulated as carpel suppression commenced in A619, but expressed at lower levels in mutant tassel primordia vs. A619 at the mid-suppression stage (Fig. 4D) (33, 34). *KIRA1* acts to regulate programmed cell death in stigmatic papillae (33). ANAC087, with ANAC046, initiates programmed cell death in the root cap, and regulates chromatin degradation following programmed cell death (34). Two proteases show a similar pattern of regulation: a cysteine protease (GRMZM2G456217) with an arabidopsis homolog (CEP1) with roles in programmed cell death of xylem elements and the tapetum (35, 36), and an uncharacterized serine protease (GRMZM2G313321, 37). These and other examples (Table 1) are consistent with roles for *GT1* and *RA3* in carpel suppression, and further point to genes that may be directly or indirectly regulated by GT1 and RA3.

**Table 1.** Putative carpel suppression genes that have been functionally characterized^1^.

| **Maize Gene ID<sup>2</sup>** | **Arabidopsis Best Hit** | **Functional Annotation** | **Biological Function** | **Citation** |
| --- | --- | --- | --- | --- |
| GRMZM2G456217 | AT5G50260 | Cysteine protease | Programmed cell death | (35, 36) |
| GRMZM2G081930 | AT4G28530 (KIRA1) | NAC domain protein 21/22 | Programmed cell death | (33) |
| **GRMZM2G181605** | AT5G18270 (ANAC087) | NAC domain protein | Programmed cell death | (34) |
| GRMZM2G313321 | AT5G36210 | serine protease | proteolysis | (37) |
| GRMZM2G085243 | AT1G16470 (PAB1) | proteasome subunit PAB1 | proteolysis |  |
| **GRMZM2G145554** | AT4G18170 (WRKY28) | WRKY DNA-binding domain protein | leaf senescence | (92) |
| **GRMZM2G104299** | AT2G33620 | AT-hook motif nuclear-localized protein 10 | ABA response; senescence (maize) | (93, 94) |
| **GRMZM5G858784** | AT1G30100 (NCED5) | nine-cis-epoxycarotenoid dioxygenase 3 | ABA biosynthesis | (95) |
| GRMZM2G052902 | AT2G24360 (RAF22) | Serine/threonine-protein kinase HT1 | ABA signaling | (96) |
| GRMZM5G884349 | AT2G35530 (bZIP16) | bZIP transcription factor superfamily protein | ABA and GA response; cell elongation | (97) |
| **GRMZM2G000264** | AT5G62670 (AHA11) | ATPase 4 plasma membrane-type | ABA response | (98) |
| GRMZM2G088299 | AT3G01490 (CBC1) | MAP kinase kinase kinase 55 | stomatal opening | (99) |
| GRMZM2G001205 | AT5G59820 (ZAT12) | ZNF5; C2H2 zinc finger protein5 | ROS response; salt and osmotic stress | (100–102) |
| **GRMZM2G152827** | AT5G14040 (PHT3;1) | mitochondrial phosphate transporter 1 | ROS accumulation; salt stress | (103, 104) |
| GRMZM5G854625 | AT1G60940 (SNRK2-10) | SNF1-related protein kinase 2.10 | ROS homeostasis; osmotic stress | (105–109) |
| **GRMZM2G080354** | AT5G10100 (TPPI) | trehalose-6-phosphate phosphatase 10 | trehalose metabolism | (110) |
| **GRMZM2G055150** | AT2G22190 (TPPE) | trpp7; trehalose-6-phosphate phosphatase 7 | trehalose metabolism | (111) |
| **GRMZM2G009223** | AT1G61800 (GPT2) | Glucose-6-phosphate/phosphate translocator | response to sucrose | (112) |
| GRMZM2G077181 | AT3G57520 (SIP2) | Galactinol--sucrose galactosyltransferase 2 | sugar metabolism | (113) |

**Table 1 (continued). Putative carpel suppression genes that have been functionally characterized<sup>1</sup>**
| **Maize Gene ID<sup>2</sup>** | **Arabidopsis Best Hit** | **Functional Annotation** | **Biological Function** | **Citation** |
| --- | --- | --- | --- | --- |
| GRMZM2G376416 | AT1G02850 (BGLU11) | Beta-glucosidase 11 | carbohydrate metabolism |  |
| **GRMZM2G170049** | AT4G25560 (LAF1) | MYB domain protein 19 | light signaling | (114, 115) |
| GRMZM2G006762 | AT5G18280 (APY2) | Putative apyrase family protein | cell elongation | (116–118) |
| GRMZM2G079957 | AT1G04240 (IAA3; SHY2) | Auxin-responsive protein IAA4 | response to auxin, cell elongation | (119–121) |
| GRMZM2G161233 | AT3G23820 (GAE6) | UDP-glucuronate 4-epimerase 1 | pectin biosynthesis; cell wall remodeling | (122) |
| GRMZM2G060824 | AT5G23630 (MIA;PDR2) | ATPase PDR2 | callose deposition; meristem maintenance | (123) |
| **GRMZM2G703858** | AT3G25070 (RIN4) | RPM1-interacting protein 4 | immunity; callose deposition | (124) |
| **GRMZM2G099454** | AT3G12500 (CHI-B) | Basic endochitinase B | immunity | (125) |
| **GRMZM2G046382** | AT1G53540 | 17.4 kDa class I heat shock protein | heat acclimation | (126) |
| GRMZM2G078025 | AT5G15450 (CLPB-P) | casein lytic proteinase B3 | heat stress; plastid development | (127, 128) |
| GRMZM2G109618 | AT3G22880 (DMC1) | DMC1; disrupted meiotic cDNA homolog1 | meiosis | (129) |

<sup>1</sup>Genes in the carpel suppression set of 73 genes with functional annotations, most of which are from arabidopsis.
<sup>2</sup>gene IDs in bold differentially expressed in *gt1* tiller buds (45). ABA = abscisic acid; GA = gibberellin; ROS = reactive oxygen species

### Trehalose levels are lower in *gt1; ra3* mutant tassels

Two *ra3* paralogs, *TPP7* (GRMZM2G080354) and *TPP10* (GRMZM2G055150), were highly upregulated during carpel suppression in A619, and strongly downregulated in *gt1; ra3* mutants (Fig. 4D, Table S3). Although RA3’s enzymatic activity is not essential in regulating meristem determinacy, it is a catalytically active TPP enzyme (22). TPP enzymes catalyze the second (and last step) of the only trehalose biosynthesis pathway in plants: they dephosphorylate T6P to produce trehalose and a free phosphate group (22, 38). The misexpression of multiple *TPP* genes led us to measure sugar and metabolite levels at the carpel suppression stage in tassel primordia (1 - 1.5 cm in length) (Fig. 4E, Fig. S3). Consistent with the downregulation of TPP genes, trehalose was substantially lower in *gt1; ra3* mutants (Fig. 4E). However, T6P levels were not concomitantly higher in mutants vs. A619. T6P is derived from the hexose phosphates glucose 6-phosphate (G6P) and UDP-glucose (UDPG), and likely signals sucrose status during plant growth and development (38– 40). Similar to T6P and most metabolites, sucrose, G6P and UDGP levels were also not significantly different between A619 and mutant tassels (Fig. 4E, Fig. S3). This indicates that while trehalose biosynthesis is impacted in *gt1; ra3* double mutants, sucrose status, as signaled by T6P (38, 40), is not affected, at least not at the level of entire tassel primordia. We reasoned that T6P levels may be unchanged because of the homeostatic control of sucrose and T6P levels (39). In support, four T6P synthases (TPSs), including at least one TPS likely to be catalytically active (41, 42), were also downregulated in *gt1; ra3* double mutants (Table S3). Taken together, these data suggest that the trehalose pathway is important in the regulation of carpel suppression.

### *ra3* enhances tillering in a *gt1* mutant background

T6P signaling has roles in regulating bud outgrowth in eudicots (43, 44), and is associated with bud outgrowth in maize (45). Notably, 24 of the 73 carpel suppression genes (~31%), including *TPP7* and *TPP10*, were differentially expressed in *gt1* vs. wild type tiller buds (45). A gene set enrichment analysis also revealed that genes differentially expressed in *gt1* vs. wild type tiller buds (45) were enriched in the differentially expressed genes in our study (Table S4), and *RA3* and *GT1* are bound and regulated by TEOSINTE BRANCHED1 (TB1), a conserved regulator of axillary bud suppression (45–47). These associations led us to ask whether *ra3* is also involved in regulating axillary bud outgrowth. We counted tillers and measured their lengths in A619, *gt1, ra3*, and *gt1; ra3* plants and found that *gt1; ra3* double mutants produced more and longer tillers than *gt1* single mutants (Fig. 5A-F). In addition, these double mutants produced more ears than single *gt1* mutants (Fig. S4). Thus, *RA3* also regulates vegetative branching in concert with *GT1*, and adds another developmental context in which both genes act to suppress growth.

**Figure 5.**
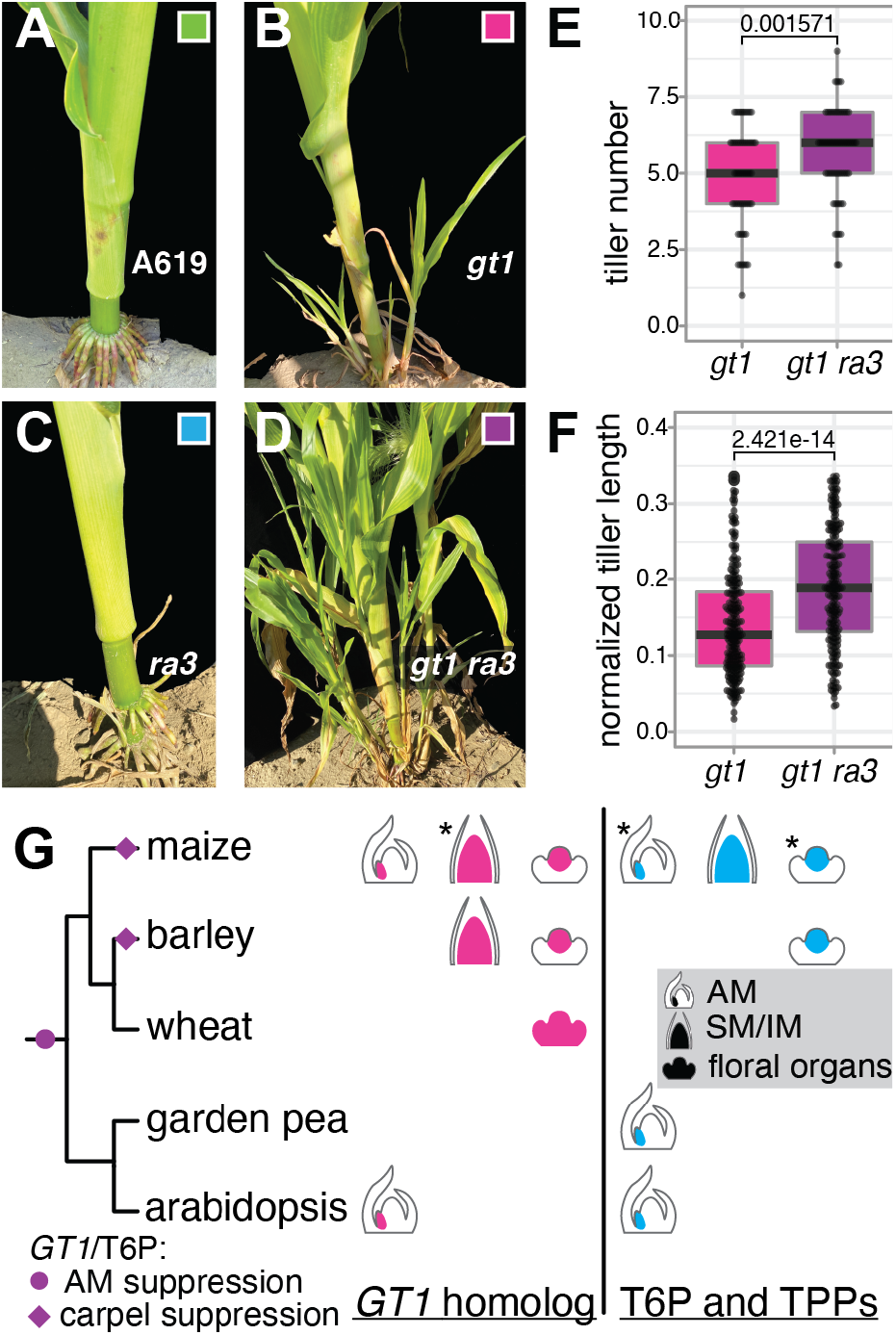
*GT1* and *RA3* act in concert to regulate carpel and axillary meristem suppression. (**A-F**) tillering was enhanced in *gt1; ra3* double mutants (p-values from ANOVA). (**G**) *GT1*-like gene and trehalose pathway-mediated regulation of axillary meristem (AM) growth has been recruited to suppress carpel primordia. Asterisks = roles for *GT1* and *RA3* we show here; citations for roles of TPP genes and T6P in the text. AM = axillary meristem, IM = inflorescence meristem, SM = spikelet meristem.

## Discussion

Organ repression is an important driving force in the evolution of floral diversity (1, 2). Here we sought to identify the genes that regulate growth suppression in maize carpels, and unexpectedly found the classic meristem determinacy gene, *RA3* (18). We show that *RA3* acts with *GT1* in multiple developmental contexts, to regulate carpel suppression, meristem determinacy, and vegetative branching. Although it is catalytically active, RA3’s enzymatic activity is not essential for regulating meristem determinacy (22). RA3 colocalizes with the transcriptional machinery in nuclear speckles in young ear primordia (22, 48). These data suggested that RA3 has an alternate ‘moonlighting’ role in regulating transcription, potentially connected to T6P signaling (22, 48). Notably, we found a similar pattern of nuclear localization for both RA3 and GT1 in carpel primordia (Fig. 4A), suggesting RA3’s moonlighting role extends to carpel suppression. Although the levels of T6P and its precursors were not significantly different between wild type and mutant tassels, trehalose levels were lower in *gt1; ra3* tassels, consistent with the downregulation of other *TPP* genes (Fig. 4D-E). These data suggest that the trehalose pathway, potentially connected to RA3’s moonlighting role, has roles in regulating carpel suppression, as it does in a number of developmental processes (40).

The trehalose pathway and *GT1-*like genes have ancient roles in regulating vegetative branching. GT1*-*like transcription factors regulate axillary meristem growth suppression in both maize and arabidopsis (17, 49). In addition, high T6P and sucrose levels lead to increased axillary bud outgrowth in arabidopsis and *Pisum sativum* (garden pea) (40, 43). In maize, high T6P levels are correlated with axillary bud outgrowth in *gt1* and *tb1* mutants (45). Notably, we found that two independent mutations in *RA3* enhance *gt1* vegetative branching (Fig. 5, Fig. S4). Taken together, these data indicate that growth regulation by the trehalose pathway and *GT1-*like genes appeared before the divergence of monocots and eudicots (Fig. 5G).

In contrast to this ancient role in vegetative branching, carpel suppression arose later and repeatedly in the grasses; for example, in the lineages leading to maize and barley (*Hordeum vulgare*) (3, 50, 51). Strikingly, carpel suppression in barley is also mediated by a *GT1* paralog, *SIX-ROWED SPIKE1* (*VRS1*), and a *RA3* paralog is downregulated in barley carpel suppression mutants (51–56). Taken together, these data suggest that *RA3* and *GT1* homologs were recruited multiple times to mediate grass carpel suppression (Fig. 5G). Furthermore, in the eudicots, *GT1-*like genes have been independently recruited to mediate unisexual flower development in *Diospyros kaki* (persimmon, 57), and a *TPP* gene occurs within the female-determining region of *Vitis vinifera* (grapevine, 58). As in the grasses, floral unisexuality arose independently in the eudicot lineages leading to persimmon and grapevine, which are separated from each other by ~120 million years, and from the grasses by ~160 million years of evolution (59). This striking convergent evolution suggests that *GT1-*like genes and TPPs were repeatedly deployed to mediate growth repression in the development of unisexual flowers.

In conclusion, we show that *GT1*-like and *RA3*-like genes control an ancient growth suppression program recruited to multiple developmental contexts. This reflects the iterative nature of plant development, where most aerial organs are leaf homologs, produced by a shoot apical meristem established early in embryogenesis (60–62). This deep homology of plant organs suggests that there may be additional cases of developmental genes seemingly functioning in discrete contexts, but whose pleiotropy is masked. Indeed, thorough genetic and phenotypic analyses continue to expose pleiotropy of developmental regulators (63–67). Recent examples include *WUSCHEL-like HOMEOBOX9 (WOX9)*, which controls both inflorescence branching and embryo development (66), and *THORN IDENTITY1*, which controls thorn development and branching in citrus (67). Detailed dissection of gene function - alone, in the context of other genes, and in different developmental contexts - is likely to reveal that this pleiotropy is widespread.

## Materials and Methods

### Plant material, growth conditions, and phenotyping

We used the *gt1-1* allele (17) for all our experiments with *gt1. gt1-1* arose in A619 and was backcrossed five times with B73 to generate the B73 introgression lines used for BSA-Seq, and for phenotypic characterizations of the *gt1; ra3* double mutants in B73 (Fig. 2C-H, Fig. S4). The *ra3* alleles used here were either those that arose in our enhancer screen (*ra3-rzl3, ra3-rzl4*) in A619, or *ra3-fea1* in B73 (18).

Plants for the *ra3-rzl4* BSA-seq, *gt1-1; ra3-rzl3* tassel and ear mature phenotypes, and *gt1-1; ra3* tiller quantification were grown at the University of Massachusetts Amherst Crop and Animal Research and Education Farm in South Deerfield, MA (~42°29’N, 72°35’W). Plants for the *gt1-1; ra3-fea1* tassel and ear branch counts were grown over 2 seasons at the UMass Amherst Crop and Animal Research and Education Farm and near Valle de Banderas, Mexico (~20°47’N, 105°15’W). Plants for SEM, metabolite measurements, and RNA-seq were grown in the College of Natural Sciences and Education Greenhouse on the UMass Amherst campus under long day conditions (16 hours light, 8 hours dark) at 28°C.

Flowers and spikelets for phenotyping were selected from the central sections of ears, or from the central sections of the tassel central spike. Tillers were measured in three planting blocks over the course of two field seasons. To account for differences in plant height, we normalized tiller length measurements to the height of the main culm.

### *rzl-4* BSA-seq

We performed an EMS mutagenesis screen as in (Neuffer et al. 1997). Briefly, *gt1-1* (A619) pollen was mutagenized with EMS and crossed onto *gt1-1* (B73) ears. M1 progeny were selfed to generate M2 families that were screened for enhanced silk growth in tassels. We identified *rzl-4* and *rzl-3* in this screen.

To map *rzl-4*, we crossed *gt1-1; rzl-4* (A619) individuals to *gt1-1* (B73) individuals, and selfed the F1 progeny to generate a F2 mapping population. In the F2 population, leaves from 238 *gt1-1; rzl-4* mutant individuals were selected for DNA extraction and pooling for bulk segregant analysis coupled to whole genome sequencing (BSA-seq) as previously described (19). Extracted genomic DNA was sequenced on an Illumina HiSeq 2500 (paired-end reads, 150 bp) at Brigham Young University.

Bioinformatic tools on the Galaxy platform were used to assess read quality, align reads, and call SNPs and indels. We used FastQC (v. 0.69) to assess read quality and we used a PHRED cutoff of 20 (68). Reads were aligned to version 3 of the B73 reference genome using Bowtie2 (v. 2.3.2.2) (69, 70). Variants were called using Samtools Mpileup (v. 2.1.3) and filtered using Varscan (v. 0.1) and Samtools filter pileup for SNPs and indels (71). Homozygous SNP variants were plotted using the R package ggplot2 (72). Candidate SNPs were identified using SnpEff (v. 4.3a) (73) and SNPs of moderate effect were filtered using Provean (21).

### Genotyping assays

*gt1-1* was genotyped with a Cleaved Amplified Polymorphic Sequences BsaJI restriction site that overlaps the *gt1-1* lesion (17, 74), resulting in cleavage of the wild-type but not mutant allele. A 287 bp fragment surrounding the *gt1-1* lesion was amplified with NEB Phusion High-Fidelity DNA polymerase with GC buffer. PCR conditions were 98 °C for 30 s, 72 °C for 15 s and 72 °C for 30 s (40 cycles). BsaJI cuts the 287 bp fragment in the wild type allele to 33 base pairs and 254 base pairs, but does not cut the mutant *gt1-1* allele.

*ra3-rzl3* was genotyped with a Derived Cleaved Amplified Polymorphic Sequences (dCAPS) Sau96I restriction site designed with dCAPS Finder 2.0 (75) (Table S5). A 119 base pair fragment surrounding the *ra3-rzl3* lesion was amplified with NEB Taq Polymerase with standard NEB buffer. PCR conditions were 95 °C for 45 s, 60 °C for 30 s and 68 °C for 30 s (40 cycles). Sau96I cuts the 119 base pair fragment in the wild type allele to 24 base pairs and 95 base pairs, but does not cut the mutant *ra3-rzl3* allele. PCR products were digested with BsaJI at 55°C for 1 hour (*gt1-1)* or Sau96I at 37°C for 1 hour (*ra3-rzl3)* and run on a 3.5% agarose gels (120V, for 60 minutes) to separate the digested bands.

### Protein Modeling

Protein homology modeling was performed with SWISS-MODEL (76). A sequence of 363 amino acids encoded by the primary *RA3* transcript was used as input into the homology search. RA3 had highest homology to the structure “*Aspergillus fumigatus* trehalose-6-phosphate phosphatase crystal form 1” (33.47%, Protein Data Bank code: 5dxl, X-ray, 1.6Å) (77). POLYVIEW-3d was used to visualize RA3 homology models with *ra3-rzl4 (78)*. ConSurf was used to assess amino acid conservation (79).

### SEMs

Scanning electron microscopy (SEM) was performed with a JEOL JCM-6000Plus Neoscope Benchtop Scanning Electron Microscope with fresh tissue samples. Tassels and ears were dissected from the main culm, their bases painted with colloidal silver paint (Ted Pella, Inc.), and placed onto pre-chilled SEM stubs with EM conductive carbon double-sided tape (Nisshin). Stubs were pre-chilled on ice for at least 15 minutes. Images were recorded under high vacuum and 5 kV voltage within 15 minutes of being in the SEM.

### RNA-seq library preparation and sequencing

We prepared a total of 32 pools of three tassels per pool at two developmental stages (1-1.1 cm and 1.3-2.0 cm) for A619, *gt1, ra3-rzl3*, and *gt1 ra3-rzl3* plants for RNA-seq. Pools for *gt1* and *ra3-rzl3* samples contained a mix of individuals either homozygous wild type or heterozygous at *ra3-rzl3* or *gt1,* respectively. Total RNA was extracted from 96 individual tassels using Trizol and the Qiagen RNeasy Plant Mini kit protocol. Tissue was collected in 1.7 mL RNase-free safe-lock tubes with ceramic grinding beads and immediately placed into liquid nitrogen. Samples were ground in a Qiagen TissueLyser II for 30 seconds and 1.0 ml of Trizol was added to each sample and mixed well by vortexing until all sample powder was thoroughly mixed. Samples were incubated for five minutes at room temperature (RT), with frequent vortexing. 0.2 ml of chloroform was added to each tube and vortexed for 15 seconds, followed by a 1 minute incubation at room temperature, and another 15 second vortex. Samples were centrifuged at 15,000 x g for 10 minutes to separate phases. After centrifugation, 200 μl of the aqueous layer was removed and added to 700 μl of Qiagen RLT buffer in a new tube. RLT buffer was prepared by adding 10 μl of β-mercaptoethanol per 1 mL RLT buffer. 500 μl of 96-100% (v/v) ethanol was then added to the 200 μl of sample now combined with 700 μl RLT buffer. Samples were mixed well by vortexing and half of the sample (~700 μl) was applied to a Qiagen MinElute pink spin column. From this point, samples were processed using the Qiagen RNeasy Plant Mini kit protocol, according to the manufacturer’s instructions. RNA quality was assessed on a 1% agarose gel run for 20 minutes at 120V and quantified using a Nanodrop 2000 spectrophotometer. Four pools were generated for each genotype and developmental stage by combining 1 ug of RNA from each of the three tassels. Total RNA from each sample was treated on-column with DNase I (NEB) for 15 minutes. RNA pools were sent to Novogene for library construction and 20 Mbp Illumina paired end 150 bp sequencing for each pool.

### RNA-seq data analysis

RNA-seq sequencing libraries were trimmed using Trimmomatic (v. 0.36.3) (80). These were then aligned to the *Zea mays* B73 v3 genome (69) and processed into BAM files using STAR (v. 2.7.0) (81). Read counts were generated with the Rsubread package function featureCounts in R (82, 83). edgeR was used to construct PCA plots of libraries (32). Because two of these libraries did not cluster with their corresponding replicates, we removed these libraries from further analysis (Table S6).

Pairwise differential expression was calculated between wild-type and each mutant using edgeR (32). Gene Ontology (GO) term enrichment was called using topGO (84) and maize-GAMER gene annotation (85). Enrichment of differentially expressed genes between wild type and *gt1* from Dong et al. 2019) in this study was assessed via Gene Set Enrichment Analysis with 1,000 permutations of the gene set in all cases except for the *gt1* vs A619 post comparison, which used 10,000 permutations (86).

### Antibody purification

GT1 (guinea pig) and RA3 (rabbit) antisera were used for antibody affinity purification to the C-terminal region of GT1 or to the N-terminal region of RA3 recombinant protein, using magnetic beads (Invitrogen), as described in (87). Validation of antibody was carried out by immunoblot following the protocol in (88), using total protein extract from wild type and *gt1* or *ra3* mutants as negative controls.

### Immunolocalizations

To perform triple whole-mount immunolocalizations, we used previously published protocols with minor modifications (48, 89). 1-1.5 cm tassels from *ra3-fea1* (B73), *gt1-1* (B73) and B73 were collected and pre-fixed at 4°C in 4% (w/v) paraformaldehyde and 2% (v/v) Tween-20 in phosphate buffered saline (PBS) for one hour, embedded in 6% agarose, sectioned at 75 μm using a vibratome (Leica), and collected in fixative solution for 2 hours more. Sections were washed and permeabilized by cell wall digestion (1% Driselase, Sigma-Aldrich; 0.5% cellulase, Sigma-Aldrich; 0.75% Pectolyase Y-23, Duchefa Biochemie) for 12 minutes at RT. Tissue was rinsed and incubated for 2 h in PBS, 2% Tween-20, rinsed 2 times and blocked with 4% (w/v) bovine serum albumin (Sigma) for 1h. The blocking solution was removed, and the tissue sections were incubated overnight at 4°C with anti-RA3 (1:200), anti-GT1 (1:75) and anti-YSPTSPS repeat S2Pho (RNA Pol II, B1 subunit; 1:200, Diagenode). The samples were washed for 8h at 4°C with gentle agitation with PBS (0.2% v/v Tween-20) replacing the buffer every 2 hours and incubated overnight at 4°C with the appropriate secondary antibodies (ThermoFisher Scientific), anti-Rabbit-Alexa 488 (RA3), anti-Mouse-Alexa 568 (RNA POL II) and anti-Guinea Pig-Alexa 647 (GT1), counterstained with DAPI (Sigma), and mounted with ProLong Gold (ThermoFisher Scientific). Immunolocalizations were repeated at least 3 times for each genotype. Images were acquired using a Zeiss LSM 780 confocal microscope. Sugars and sugar alcohols were measured by LC-MS/MS as described by (90).

### Metabolite measurements

Tassels between 1-1.4 cm were dissected from A619, *gt1-1, ra3-rzl3; gt1-1/+*, and *gt1-1; ra3-rzl3* individuals and immediately frozen in liquid nitrogen. Tassels of all four genotypes were sampled in four batches over a four week period. All individuals in a batch were sampled at 15h00 to minimize variability between batches. T6P and sugar metabolites were extracted from single tassels using a protocol from (39). T6P, other phosphorylated intermediates, and organic acids were measured by anion-exchange high performance liquid chromatography coupled to tandem mass spectrometry (LC-MS/MS) as described by (39) with modifications as described by (91).

## Data Availability

Raw sequencing data is available at NCBI BioProjects (RNA-seq: PRJNA657042; *ra3-rzl4* BSA-seq: PRJNA656888).

## Acknowledgements

We thank members of the Bartlett lab for thoughtful discussion and comments on this work, and ‘critical friends’, Zachary Lippman and Beth Thompson. We thank Amber De Neve, Ed Wilcox, Jae Hyung Lee, and Xiaosa Xu for help with experimentation, and Dan Jones, Chris Joyner, Chris Phillips, and Neal Woodard, for essential help with nursery management and plant care. This work was supported by the National Science Foundation (IOS-1652380 to M.B., IOS-1755141 to D.J.), the USDA National Institute of Food and Agriculture (NIFA-2019-67012-29654 to J.P.G.), the University of Massachusetts, and the Max Planck Societ

## Notes

### Competing Interest Statement

The authors have declared no competing interest.

## References

1. L. Ronse De Craene, Understanding the role of floral development in the evolution of angiosperm flowers: clarifications from a historical and physico-dynamic perspective. J. Plant Res. 131, 367–393 (2018).

2. P. J. Rudall, R. M. Bateman, Evolution of zygomorphy in monocot flowers: Iterative patterns and developmental constraints. New Phytol. 162, 25–44 (2004).

3. E. A. Kellogg, Flowering Plants. Monocots: Poaceae (Springer International Publishing, 2015).

4. S. Sakuma, T. Schnurbusch, Of floral fortune: tinkering with the grain yield potential of cereal crops. New Phytol. 225, 1873–1882 (2020).

5. J. Friedman, S. C. H. Barrett, Wind of change: new insights on the ecology and evolution of pollination and mating in wind-pollinated plants. Ann. Bot. 103, 1515–1527 (2009).

6. L. Chen, Y.-G. Liu, Male sterility and fertility restoration in crops. Annu. Rev. Plant Biol. 65, 579–606 (2014).

7. P. C. Cheng, R. I. Greyson, D. B. Walden, Organ initiation and the development of unisexual flowers in the tassel and ear of Zea mays. Am. J. Bot. 70, 450–462 (1983).

8. A. Calderon-Urrea, S. L. Dellaporta, Cell death and cell protection genes determine the fate of pistils in maize. Development 126, 435–441 (1999).

9. A. DeLong, A. Calderon-Urrea, S. L. Dellaporta, Sex determination gene TASSELSEED2 of maize encodes a short-chain alcohol dehydrogenase required for stage-specific floral organ abortion. Cell 74, 757–768 (1993).

10. I. F. Acosta, et al., tasselseed1 is a lipoxygenase affecting jasmonic acid signaling in sex determination of maize. Science 323, 262–265 (2009).

11. N. B. Best, et al., nana plant2 encodes a maize ortholog of the Arabidopsis brassinosteroid biosynthesis gene DWARF1, identifying developmental interactions between brassinosteroids and gibberellins. Plant Physiol. 171, 2633–2647 (2016).

12. G. Chuck, R. Meeley, E. Irish, H. Sakai, S. Hake, The maize tasselseed4 microRNA controls sex determination and meristem cell fate by targeting Tasselseed6/indeterminate spikelet1. Nat. Genet. 39, 1517–1521 (2007).

13. C. Lunde, A. Kimberlin, S. Leiboff, A. J. Koo, S. Hake, Tasselseed5 overexpresses a wound-inducible enzyme, ZmCYP94B1, that affects jasmonate catabolism, sex determination, and plant architecture in maize. Commun Biol 2, 114 (2019).

14. G. Chuck, R. Meeley, S. Hake, Floral meristem initiation and meristem cell fate are regulated by the maize AP2 genes ids1 and sid1. Development 135, 3013–3019 (2008).

15. S. E. Parkinson, S. M. Gross, J. B. Hollick, Maize sex determination and abaxial leaf fates are canalized by a factor that maintains repressed epigenetic states. Dev. Biol. 308, 462–473 (2007).

16. N. H. Nickerson, E. E. Dale, Tassel modifications in Zea mays. Ann. Mo. Bot. Gard. 42, 195–211 (1955).

17. C. J. Whipple, et al., grassy tillers1 promotes apical dominance in maize and responds to shade signals in the grasses. Proc. Natl Acad. Sci. USA 108, E506–E512 (2011).

18. N. Satoh-Nagasawa, N. Nagasawa, S. Malcomber, H. Sakai, D. Jackson, A trehalose metabolic enzyme controls inflorescence architecture in maize. Nature 441, 227 (2006).

19. H. Klein, et al., Bulked-segregant analysis coupled to whole genome sequencing (BSA-Seq) for rapid gene cloning in maize. G3 8, 3583–3592 (2018).

20. Z. Liang, J. C. Schnable, RNA-Seq based analysis of population structure within the maize inbred B73. PLoS One 11, e0157942 (2016).

21. Y. Choi, A. P. Chan, PROVEAN web server: a tool to predict the functional effect of amino acid substitutions and indels. Bioinformatics 31, 2745–2747 (2015).

22. H. Claeys, et al., Control of meristem determinacy by trehalose 6-phosphate phosphatases is uncoupled from enzymatic activity. Nat Plants 5, 352–357 (2019).

23. E. E. Irish, T. Nelson, Sex determination in monoecious and dioecious plants. Plant Cell 1, 737–744 (1989).

24. J. C. Kim, et al., Cell cycle arrest of stamen initials in maize sex determination. Genetics 177, 2547–2551 (2007).

25. L. Colombo, G. Marziani, S. Masiero, P. E. Wittich, BRANCHED SILKLESS mediates the transition from spikelet to floral meristem during Zea mays ear development. Plant J. 16, 355–363 (1998).

26. J. Strable, E. Vollbrecht, Maize YABBY genes drooping leaf1 and drooping leaf2 regulate floret development and floral meristem determinacy. Development 146, dev171181 (2019).

27. B. E. Thompson, et al., bearded-ear Encodes a MADS Box transcription factor critical for maize floral development. Plant Cell 21, 2578–2590 (2009).

28. M. Mena, et al., Diversification of C-function activity in maize flower development. Science 274, 1537–1540 (1996).

29. E. Irish, Class II tassel seed mutations provide evidence for multiple types of inflorescence meristems in maize (Poaceae). Am. J. Bot. 84, 1502 (1997).

30. R. Reinheimer, F. O. Zuloaga, A. C. Vegetti, R. Pozner, Diversification of inflorescence development in the PCK clade (Poaceae: Panicoideae: Paniceae). Am. J. Bot. 96, 549–564 (2009).

31. C. J. Whipple, Grass inflorescence architecture and evolution: the origin of novel signaling centers. New Phytol. 216, 367–372 (2017).

32. M. D. Robinson, D. J. McCarthy, G. K. Smyth, edgeR: a Bioconductor package for differential expression analysis of digital gene expression data. Bioinformatics 26, 139–140 (2010).

33. Z. Gao, et al., KIRA1 and ORESARA1 terminate flower receptivity by promoting cell death in the stigma of Arabidopsis. Nature Plants 4, 365–375 (2018).

34. M. Huysmans, et al., NAC transcription factors ANAC087 and ANAC046 control distinct aspects of programmed cell death in the Arabidopsis columella and lateral root cap. Plant Cell 30, 2197–2213 (2018).

35. J. Han, et al., The papain-like cysteine protease CEP1 is involved in programmed cell death and secondary wall thickening during xylem development in Arabidopsis. J. Exp. Bot. 70, 205–215 (2019).

36. D. Zhang, et al., The cysteine protease CEP1, a key executor involved in tapetal programmed cell death, regulates pollen development in Arabidopsis. Plant Cell 26, 2939–2961 (2014).

37. A. Tsuji, Y. Fujisawa, T. Mino, K. Yuasa, Identification of a plant aminopeptidase with preference for aromatic amino acid residues as a novel member of the prolyl oligopeptidase family of serine proteases. J. Biochem. 150, 525–534 (2011).

38. M. J. Paul, L. F. Primavesi, D. Jhurreea, Y. Zhang, Trehalose metabolism and signaling. Annu. Rev. Plant Biol. 59, 417–441 (2008).

39. J. E. Lunn, et al., Sugar-induced increases in trehalose 6-phosphate are correlated with redox activation of ADPglucose pyrophosphorylase and higher rates of starch synthesis in Arabidopsis thaliana. Biochem. J 397, 139–148 (2006).

40. F. Fichtner, J. E. Lunn, The role of trehalose 6-phosphate (Tre6P) in plant metabolism and development. Annu. Rev. Plant Biol. 72, 737–760 (2021).

41. M.-L. Zhou, et al., Trehalose metabolism-related genes in maize. J. Plant Growth Regul. 33, 256–271 (2014).

42. C. Henry, et al., The trehalose pathway in maize: conservation and gene regulation in response to the diurnal cycle and extended darkness. J. Exp. Bot. 65, 5959–5973 (2014).

43. F. Fichtner, et al., Trehalose 6-phosphate is involved in triggering axillary bud outgrowth in garden pea (Pisum sativum L.). Plant J. 92, 611–623 (2017).

44. C. Tarancón, E. González-Grandío, J. C. Oliveros, M. Nicolas, P. Cubas, A conserved carbon starvation response underlies bud dormancy in woody and herbaceous species. Front. Plant Sci. 8, 788 (2017).

45. Z. Dong, et al., The regulatory landscape of a core maize domestication module controlling bud dormancy and growth repression. Nat. Commun. 10, 3810 (2019).

46. M. Wang, et al., BRANCHED1: A Key Hub of Shoot Branching. Front. Plant Sci. 10, 76 (2019).

47. J. Doebley, A. Stec, L. Hubbard, The evolution of apical dominance in maize. Nature 386, 485–488 (1997).

48. E. Demesa-Arevalo, M. J. Abraham-Juarez, X. Xu, Maize RAMOSA3 accumulates in nuclear condensates enriched in RNA POLYMERASE II isoforms during the establishment of axillary meristem determinacy. bioRxiv (2021).

49. E. González-Grandío, et al., Abscisic acid signaling is controlled by a BRANCHED1/HD-ZIP I cascade in Arabidopsis axillary buds. Proc. Natl. Acad. Sci. USA 114, E245–E254 (2017).

50. L. G. Le Roux, E. A. Kellogg, Floral development and the formation of unisexual spikelets in the Andropogoneae (Poaceae). Am. J. Bot. 86, 354–366 (1999).

51. R. Koppolu, et al., Six-rowed spike4 (Vrs4) controls spikelet determinacy and row-type in barley. Proc. Natl. Acad. Sci. USA 110, 13198–13203 (2013).

52. Y. Shang, et al., A CYC/TB1-type TCP transcription factor controls spikelet meristem identity in barley. J. Exp. Bot. 71, 7118–7131 (2020).

53. N. Poursarebani, et al., COMPOSITUM 1 contributes to the architectural simplification of barley inflorescence via meristem identity signals. Nat. Commun. 11, 5138 (2020).

54. J. Thiel, et al., Transcriptional landscapes of floral meristems in barley. Sci Adv 7, eabf0832 (2021).

55. S. Sakuma, et al., Divergence of expression pattern contributed to neofunctionalization of duplicated HD-Zip I transcription factor in barley. New Phytol. 197, 939–948 (2013).

56. M. Zwirek, R. Waugh, S. M. McKim, Interaction between row-type genes in barley controls meristem determinacy and reveals novel routes to improved grain. New Phytol. 221, 1950–1965 (2019).

57. T. Akagi, I. M. Henry, R. Tao, L. Comai, A Y-chromosome–encoded small RNA acts as a sex determinant in persimmons. Science 346, 646–650 (2014).

58. C. Zou, et al., Multiple independent recombinations led to hermaphroditism in grapevine. Proc. Natl. Acad. Sci. USA 118, e2023548118 (2021).

59. S. Kumar, G. Stecher, M. Suleski, S. B. Hedges, TimeTree: A resource for timelines, timetrees, and divergence times. Mol. Biol. Evol. 34, 1812–1819 (2017).

60. D. R. Kaplan, T. J. Cooke, Fundamental concepts in the embryogenesis of dicotyledons: A morphological interpretation of embryo mutants. Plant Cell 9, 1903–1919 (1997).

61. S. Leiboff, S. Hake, Reconstructing the transcriptional ontogeny of maize and sorghum supports an inverse hourglass model of inflorescence development. Curr. Biol. 29, 3410–3419.e3 (2019).

62. J. W. Von Goethe, G. L. Miller, The metamorphosis of plants (MIT Press Cambridge, 2009).

63. S. A. Kessler, et al., Conserved molecular components for pollen tube reception and fungal invasion. Science 330, 968–971 (2010).

64. C. Li, et al., Glycosylphosphatidylinositol-anchored proteins as chaperones and co-receptors for FERONIA receptor kinase signaling in Arabidopsis. Elife 4, e06587 (2015).

65. T. Rosas-Diaz, et al., A virus-targeted plant receptor-like kinase promotes cell-to-cell spread of RNAi. Proc. Natl. Acad. Sci. USA 115, 1388–1393 (2018).

66. A. Hendelman, et al., Conserved pleiotropy of an ancient plant homeobox gene uncovered by cis-regulatory dissection. Cell 184, 1724–1739.e16 (2021).

67. F. Zhang, et al., Reprogramming of stem cell activity to convert thorns into branches. Curr. Biol. 30, 2951–2961.e5 (2020).

68. S. Andrews, FastQC: a quality control tool for high throughput sequence data. Available online at: http://www.bioinformatics.babraham.ac.uk/projects/fastqc/ (2010).

69. P. S. Schnable, et al., The B73 maize genome: complexity, diversity, and dynamics. Science 326, 1112–1115 (2009).

70. B. Langmead, C. Trapnell, M. Pop, S. Salzberg, Ultrafast and memory-efficient alignment ofshort DNA sequences to the human genome. Genome Biol. 10, R25 (2009).

71. H. Li, et al., The sequence alignment/map format and SAMtools. Bioinformatics 25, 2078–2079 (2009).

72. H. Wickham, ggplot2: Elegant Graphics for Data Analysis (Springer, 2016).

73. P. Cingolani, et al., A program for annotating and predicting the effects of single nucleotide polymorphisms, SnpEff: SNPs in the genome of Drosophila melanogaster strain w1118; iso-2; iso-3. Fly 6, 80–92 (2012).

74. G. P. Rédei, Ed., “CAPS (cleaved amplified polymorphic sequences)” in Encyclopedia of Genetics, Genomics, Proteomics and Informatics, (Springer Netherlands, 2008), pp. 270–270.

75. M. M. Neff, E. Turk, M. Kalishman, Web-based primer design for single nucleotide polymorphism analysis. Trends Genet. 18, 613–615 (2002).

76. A. Waterhouse, et al., SWISS-MODEL: homology modelling of protein structures and complexes. Nucleic Acids Res. 46, W296–W303 (2018).

77. Y. Miao, et al., Structures of trehalose-6-phosphate phosphatase from pathogenic fungi reveal the mechanisms of substrate recognition and catalysis. Proc. Natl. Acad. Sci. USA 113, 7148–7153 (2016).

78. A. Porollo, J. Meller, Versatile annotation and publication quality visualization of protein complexes using POLYVIEW-3D. BMC Bioinformatics 8, 316 (2007).

79. H. Ashkenazy, et al., ConSurf 2016: an improved methodology to estimate and visualize evolutionary conservation in macromolecules. Nucleic Acids Res. 44, W344–50 (2016).

80. A. M. Bolger, M. Lohse, B. Usadel, Trimmomatic: a flexible trimmer for Illumina sequence data. Bioinformatics 30, 2114–2120 (2014).

81. A. Dobin, et al., STAR: ultrafast universal RNA-seq aligner. Bioinformatics 29, 15–21 (2013).

82. Y. Liao, G. K. Smyth, W. Shi, The R package Rsubread is easier, faster, cheaper and better for alignment and quantification of RNA sequencing reads. Nucleic Acids Res. 47, e47 (2019).

83. R. Ihaka, R. Gentleman, R: A language for data analysis and graphics. J. Comput. Graph. Stat. 5, 299–314 (1996).

84. A. Alexa, J. Rahnenfuhrer, topGO: Enrichment Analysis for Gene Ontology. R package version 2.44.0 (2016).

85. K. Wimalanathan, I. Friedberg, C. M. Andorf, C. J. Lawrence-Dill, Maize GO annotation - methods, evaluation, and review (maize-GAMER). Plant Direct 2, e00052 (2018).

86. A. Subramanian, et al., Gene set enrichment analysis: a knowledge-based approach for interpreting genome-wide expression profiles. Proc. Natl. Acad. Sci. USA 102, 15545–15550 (2005).

87. G. S. Chuck, P. J. Brown, R. Meeley, S. Hake, Maize SBP-box transcription factors unbranched2 and unbranched3 affect yield traits by regulating the rate of lateral primordia initiation. Proc. Natl. Acad. Sci. USA 111, 18775–18780 (2014).

88. M. J. Abraham-Juárez, Western blot in maize. Bio-101 e3257 (2019).

89. T. M. Tran, et al., An optimized whole-mount immunofluorescence method for shoot apices. Curr Protoc 1, e101 (2021).

90. R. Feil, J. E. Lunn, Quantification of soluble sugars and sugar alcohols by LC-MS/MS. Methods Mol. Biol. 1778, 87–100 (2018).

91. C. M. Figueroa, et al., Trehalose 6-phosphate coordinates organic and amino acid metabolism with carbon availability. Plant J. 85, 410–423 (2016).

92. T. Tian, et al., Arabidopsis FAR-RED ELONGATED HYPOCOTYL3 Integrates Age and Light Signals to Negatively Regulate Leaf Senescence. Plant Cell 32, 1574–1588 (2020).

93. M. M. Wong, et al., Phosphoproteomics of Arabidopsis Highly ABA-Induced1 identifies AT-Hook–Like10 phosphorylation required for stress growth regulation. Proc. Natl. Acad. Sci. USA 116, 2354–2363 (2019).

94. R. S. Sekhon, et al., Integrated genome-scale analysis identifies novel genes and networks underlying senescence in maize. Plant Cell 31, 1968–1989 (2019).

95. B.-C. Tan, et al., Molecular characterization of the Arabidopsis 9-cis epoxycarotenoid dioxygenase gene family. Plant J. 35, 44–56 (2003).

96. Y. Kamiyama, et al., Arabidopsis group C Raf-like protein kinases negatively regulate abscisic acid signaling and are direct substrates of SnRK2. Proc. Natl. Acad. Sci. USA 118, e2100073118 (2021).

97. W.-P. Hsieh, H.-L. Hsieh, S.-H. Wu, Arabidopsis bZIP16 transcription factor integrates light and hormone signaling pathways to regulate early seedling development. Plant Cell 24, 3997–4011 (2012).

98. X. Zhang, et al., Inhibition of blue light-dependent H+ pumping by abscisic acid through hydrogen peroxide-induced dephosphorylation of the plasma membrane H+-ATPase in guard cell protoplasts. Plant Physiol. 136, 4150–4158 (2004).

99. A. Hiyama, et al., Blue light and CO2 signals converge to regulate light-induced stomatal opening. Nat. Commun. 8, 1284 (2017).

100. S. Davletova, K. Schlauch, J. Coutu, R. Mittler, The zinc-finger protein Zat12 plays a central role in reactive oxygen and abiotic stress signaling in Arabidopsis. Plant Physiol. 139, 847–856 (2005).

101. I. De Clercq, et al., Integrative inference of transcriptional networks in Arabidopsis yields novel ROS signalling regulators. Nat Plants 7, 500–513 (2021).

102. L. Rizhsky, S. Davletova, H. Liang, R. Mittler, The zinc finger protein Zat12 is required for Cytosolic Ascorbate Peroxidase 1 expression during oxidative stress in Arabidopsis. Journal of Biological Chemistry 279, 11736–11743 (2004).

103. F. Jia, et al., Overexpression of mitochondrial phosphate transporter 3 severely hampers plant development through regulating mitochondrial function in Arabidopsis. PLoS One 10, e0129717 (2015).

104. W. Zhu, et al., The mitochondrial phosphate transporters modulate plant responses to salt stress via affecting ATP and gibberellin metabolism in Arabidopsis thaliana. PLoS One 7, e43530 (2012).

105. H. Fujii, P. E. Verslues, J.-K. Zhu, Arabidopsis decuple mutant reveals the importance of SnRK2 kinases in osmotic stress responses in vivo. Proc. Natl. Acad. Sci. USA 108, 1717–1722 (2011).

106. F. Soma, et al., ABA-unresponsive SnRK2 protein kinases regulate mRNA decay under osmotic stress in plants. Nat Plants 3, 16204 (2017).

107. N. Fàbregas, T. Yoshida, A. R. Fernie, Role of Raf-like kinases in SnRK2 activation and osmotic stress response in plants. Nat. Commun. 11, 6184 (2020).

108. D. Kawa, et al., SnRK2 protein kinases and mRNA decapping machinery control root development and response to salt. Plant Physiol. 182, 361–377 (2020).

109. K. P. Szymańska, L. Polkowska-Kowalczyk, M. Lichocka, J. Maszkowska, G. Dobrowolska, SNF1-related protein kinases SnRK2.4 and SnRK2.10 modulate ROS homeostasis in plant response to salt stress. Int. J. Mol. Sci. 20, 143 (2019).

110. Q. Lin, S. Wang, Y. Dao, J. Wang, K. Wang, Arabidopsis thaliana trehalose-6-phosphate phosphatase gene TPPI enhances drought tolerance by regulating stomatal apertures. J. Exp. Bot. 71, 4285–4297 (2020).

111. W. Wang, et al., Trehalose-6-phosphate phosphatase E modulates ABA-controlled root growth and stomatal movement in Arabidopsis. J. Integr. Plant Biol. 62, 1518–1534 (2020).

112. B. C. Dyson, R. E. Webster, G. N. Johnson, GPT2: a glucose 6-phosphate/phosphate translocator with a novel role in the regulation of sugar signalling during seedling development. Ann. Bot. 113, 643–652 (2014).

113. J. Ma, et al., The sucrose-regulated Arabidopsis transcription factor bZIP11 reprograms metabolism and regulates trehalose metabolism. New Phytol. 191, 733–745 (2011).

114. I.-C. Jang, R. Henriques, N.-H. Chua, Three transcription factors, HFR1, LAF1 and HY5, regulate largely independent signaling pathways downstream of phytochrome A. Plant Cell Physiol. 54, 907–916 (2013).

115. S. W. Yang, I.-C. Jang, R. Henriques, N.-H. Chua, FAR-RED ELONGATED HYPOCOTYL1 and FHY1-LIKE associate with the Arabidopsis transcription factors LAF1 and HFR1 to transmit phytochrome A signals for inhibition of hypocotyl elongation. Plant Cell 21, 1341–1359 (2009).

116. J. Wu, et al., Apyrases (nucleoside triphosphate-diphosphohydrolases) play a key role in growth control in Arabidopsis. Plant Physiol. 144, 961–975 (2007).

117. C. Wolf, M. Hennig, D. Romanovicz, I. Steinebrunner, Developmental defects and seedling lethality in apyrase AtAPY1 and AtAPY2 double knockout mutants. Plant Mol. Biol. 64, 657–672 (2007).

118. G. Clark, et al., Apyrase (nucleoside triphosphate-diphosphohydrolase) and extracellular nucleotides regulate cotton fiber elongation in cultured ovules. Plant Physiol. 152, 1073–1083 (2010).

119. Y. Xi, et al., IAA3-mediated repression of PIF proteins coordinates light and auxin signaling in Arabidopsis. PLoS Genet. 17, e1009384 (2021).

120. C. Weiste, et al., The Arabidopsis bZIP11 transcription factor links low-energy signalling to auxin-mediated control of primary root growth. PLoS Genet. 13, e1006607 (2017).

121. Q. Tian, J. W. Reed, Control of auxin-regulated root development by the Arabidopsis thaliana SHY2/IAA3 gene. Development 126, 711–721 (1999).

122. G. Bethke, et al., Pectin biosynthesis is critical for cell wall integrity and immunity in Arabidopsis thaliana. Plant Cell 28, 537–556 (2016).

123. J. Müller, et al., Iron-dependent callose deposition adjusts root meristem maintenance to phosphate availability. Dev. Cell 33, 216–230 (2015).

124. E.-H. Chung, F. El-Kasmi, Y. He, A. Loehr, J. L. Dangl, A plant phosphoswitch platform repeatedly targeted by type III effector proteins regulates the output of both tiers of plant immune receptors. Cell Host Microbe 16, 484–494 (2014).

125. L. K. Hawkins, et al., Characterization of the maize chitinase genes and their effect on Aspergillus flavus and aflatoxin accumulation resistance. PLoS One 10, e0126185 (2015).

126. N. Yamaguchi, et al., H3K27me3 demethylases alter HSP22 and HSP17.6C expression in response to recurring heat in Arabidopsis. Nat. Commun. 12, 3480 (2021).

127. I. L. Parcerisa, G. L. Rosano, E. A. Ceccarelli, Biochemical characterization of ClpB3, a chloroplastic disaggregase from Arabidopsis thaliana. Plant Mol. Biol. 104, 451–465 (2020).

128. F. Myouga, R. Motohashi, T. Kuromori, N. Nagata, K. Shinozaki, An Arabidopsis chloroplast-targeted Hsp101 homologue, APG6, has an essential role in chloroplast development as well as heat-stress response. Plant J. 48, 249–260 (2006).

129. H. Chen, et al., RAD51 supports DMC1 by inhibiting the SMC5/6 complex during meiosis. Plant Cell 33, 2869–2882 (2021).

