## Supplemental Figures and Tables for "Recruitment of an ancient branching program to suppress carpel development in maize flowers"

**This PDF file includes:**

Figures S1 to S3
Tables S1 to S5

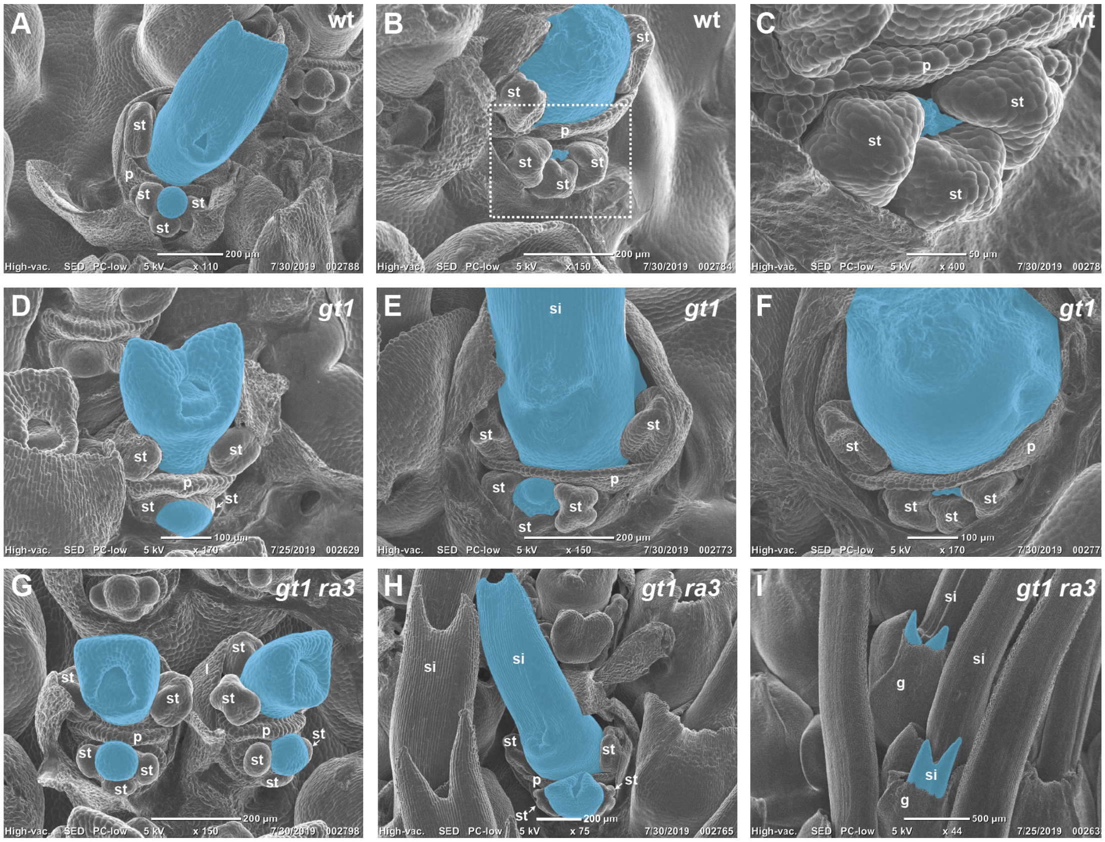


**Figure S1.** Carpel suppression was disrupted in the lower flowers of *gt1; ra3* ear spikelets. Ear flower development in (**A-C**) wild type (A619), (**D-F**) *gt1*, and (**G-I**) *gt1; ra3* double mutants. Outlined regions in B shown at higher magnification in C. Some gynoecia false-colored blue; glumes dissected off of spikelets in A-I. Silks broken off in B and F. g = glume, l = lemma, p = palea, si = silk, st = stamen primordium

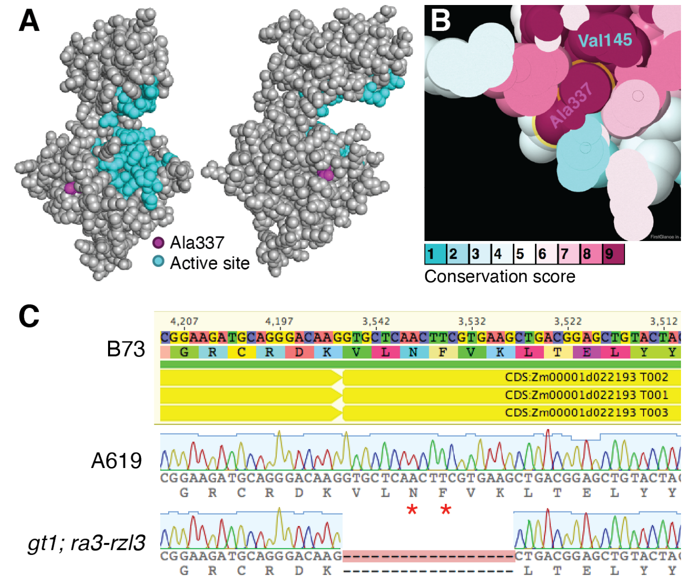


**Figure S2**. *rzl-3* and *rzl-4* mutants both harbor mutations in *RA3*. (**A-B**) A missense mutation in *ra3-rzl4* impacts an amino acid in contact with the active site. Predicted structure of RA3 protein modeled on a TPP from Aspergillus fumigatus, shown from two different views. Active site residues in teal, Ala337 in magenta (Val337 in *ra3-rzl4* gene product). (**C**) Consurf conservation scores mapped onto the modeled RA3 structure. Both Ala337 and the active site residue Val145 are deeply conserved, and contact each other in the model. (**D**) A splice site mutation in *ra3-rzl3* results in a deletion of six codons, including codons for two active site residues (red asterisks).

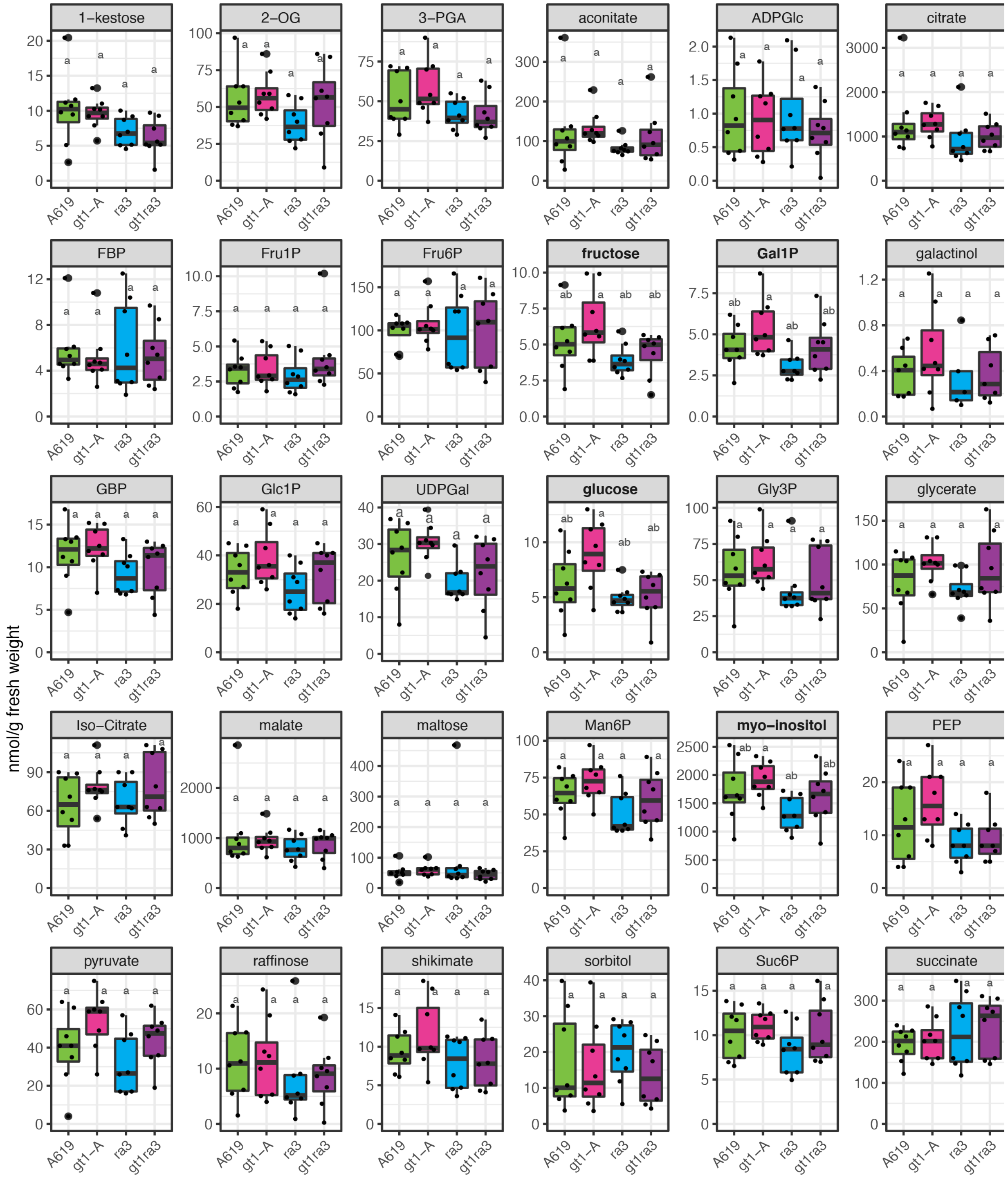


**Figure S3.** Metabolite levels in *gt1, ra3,* and *gt1; ra3* double mutant tassels were unchanged relative to wild type (A619). Different letters indicate statistically significant differences ($p < 0.05$, ANOVA, Tukey's post-hoc). 2-OG, 2-oxoglutarate; 3-PGA,3-Phosphoglyceric acid; ADPGlc, adenosine diphosphoglucose; FBP, fructose 1,6-bisphosphate; Fru1P, fructose-1-phosphate; Fru6P, fructose-6-phosphate; Gal1P, galactose-1-phosphate; G1,6BP, glucose 1,6-bisphosphate; Glc, glucose; Glc1P, glucose-1-phosphate; Glc6P, glucose-6-phosphate; Gly3P, glycerol 3-phosphate; Man6P, Mannose-6-phosphate; PEP, phosphoenolpyruvate; Suc6P, sucrose-6-phosphate; T6P, trehalose-6-phosphate; UDPGa, uridine diphosphogalactose; UDPGlc, uridine diphosphoglucose.

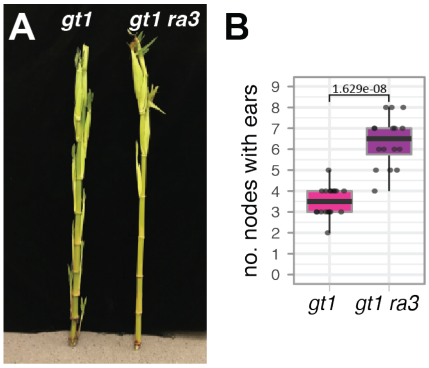


**Figure S4.** *gt1; ra3-fea1* double mutants had more axillary branches than *gt1* single mutants in the B73 genetic background. Statistical significance assessed using Student's t-test.

**Table S1.** Complementation tests between *rzl-3* and *rzl-4*.

| Generation (cross) | Progeny Phenotype |
|---|---|
| F1 (*gt1; rzl-3 X gt1; rzl4)* | 35/35 *rzl* |
| F2 (F1 selfed) | 14/14 *rzl* |

**Table S2.** Stamen and lodicule counts[1]

| Genotype (plants/flowers) | Stamens | Lodicules | Ear Branches |
|---|---|---|---|
| A619 (5/60) | 3 (0.5) | 2 (0.7) | 0 (0) |
| *gt1; ra3-rzl3* (5/60) | 3 (0.2) | 2 (0) | n.d. |
| *gt1; ra3-rzl3/+* (5/60) | 3 (0.4) | 2 (0.3) | 0 (0) |
| *ra3-rzl3; gt1/+* | n.d. | n.d. | 0 (0) |

n.d. = not determined. Means of organ and branch counts reported, standard deviations in parentheses

**Table S3.** Enrichment of differentially expressed genes from *gt1* tiller bud studies in Dong et al. (2019) within this study's gene expression dataset (q-values reported).

| | | ***gt1* vs. A619** | | ***ra3* vs. A619** | | ***gt1 ra3* vs. A619** | |
|---|---|---|---|---|---|---|---|
| **Gene Set (*gt1* vs. wild type)** | **Size** | **pre** | **mid** | **pre** | **mid** | **pre** | **mid** |
| Genes up in *gt1* tiller buds (12 DAP) | 3331 | 0.039 | 0 | 0.006 | 0.009 | 0 | 0.003 |
| Genes up in *gt1* tiller buds (8 DAP) | 308 | 0.666 | - | 0.023 | 0.038 | - | 0.018 |
| Genes up in wild type tiller buds (8 DAP) | 706 | 0.15 | - | - | 0.001 | 0 | - |
| Genes up in wild type tiller buds (12 DAP) | 3696 | - | - | - | 0.001 | - | - |

DAP = days after planting; - = no enrichment

**Table S4.** Primers and genotyping assays

| Primer Name | Sequence | PCR/Genotyping Protocol |
|---|---|---|
| umc1029 F | AACACCTGCTGGATATGGATCACT | B73 ~150 bp, A619 ~130 bp. |
| umc1029 R | GGAAGAAAAATGTCGACCTGCTC | |
| SSR6c7 172481663 F | GCTTAACAAGTAGCACAGCGC | B73 ~139 bp, A619 ~117 bp. |
| SSR6c7 172481802 R | TGTAATATCGGTCGGCGTCG | |
| SSR5c7 172712040 F | GCATTGATCCGGCAGACTCA | B73 ~139 bp, A619 ~128 bp. |
| SSR5c7 172712128 R | TATTGGCACTCTTGTCGGCC | |
| Zm00001d022259 F | TCAAGAAGAAGCTCCGGCAG | FokI digest. B73 ~700 bp, A619 ~780 bp. |
| Zm00001d022259 R | TGTGCTGGAGATGGAGTTGG | |
| umc1154 F | CCACCACAAGACAAGACAAGAATG | B73 ~100 bp, A619 ~150 bp. |
| umc1154 R | CCTGATCGATCTCATCGTCGT | |
| *ra3*-rzl3 dCAPS F | CGTCAGCTTCACGAAGTTGAGGAC | Sau96I dCAPS digest. A619 ~100 bp, *ra3-rzl3* ~120 bp. |
| *ra3*-rzl3 dCAPS R | AGTAGCACTCTGGGAAAGCG | |
| *ra3* CDS F | ATGACGAAGCACGCCGCCT | Amplifies full length *ra3* CDS with stop codon. |
| *ra3* CDS R | TCATGGTTGGCGCGCCCCCTTC | |
| *gt1*-1 F | GACGAGCAGGCCAGGAAGC | BsaJI digest. 254 bp wild type, ~ 287 bp *gt1-1*. |
| *gt1*-1 R | CGAGGTTCGCGGATCAGTTAC | |

**Table S5.** Read counts and mapped reads.

| Sample ID | Q30(%) | Trimmed Input STAR | Aligned Reads | Aligned(%) | Counted Reads | Counted(%) |
|---|---|---|---|---|---|---|
| **A619 pre 1** | 94.95 | 23812814 | 21313511 | 89.5 | 18443314 | 86.5 |
| **A619 pre 2** | 94.81 | 35390356 | 28623936 | 80.9 | 24641460 | 86.1 |
| **A619 pre 3** | 94.75 | 28222606 | 25439732 | 90.1 | 22289399 | 87.6 |
| **A619 pre 4** | 94.78 | 25605371 | 23126536 | 90.3 | 20231314 | 87.5 |
| ***gt1* pre 1** | 94.9 | 32147599 | 25368882 | 78.9 | 21758736 | 85.8 |
| ***gt1* pre 2** | 94.77 | 27801802 | 24988706 | 89.9 | 21708581 | 86.9 |
| ***gt1* pre 3*** | 94.97 | 25864580 | 23034911 | 89.1 | 19806171 | 86.0 |
| ***gt1* pre 4** | 94.74 | 29572385 | 22971081 | 77.7 | 19504052 | 84.9 |
| ***ra3* pre 1** | 94.69 | 26941203 | 24232410 | 89.9 | 21129352 | 87.2 |
| ***ra3* pre 2** | 94.88 | 24118934 | 21731014 | 90.1 | 18964603 | 87.3 |
| ***ra3* pre 3** | 94.97 | 24631267 | 22202776 | 90.1 | 19532018 | 88.0 |
| ***ra3* pre 4** | 94.9 | 30210805 | 26984830 | 89.3 | 23650751 | 87.6 |
| ***gt1 ra3* pre 1** | 94.35 | 25678998 | 23032722 | 89.7 | 20139067 | 87.4 |
| ***gt1 ra3* pre 2** | 94.74 | 28418191 | 25539756 | 89.9 | 22216884 | 87.0 |
| ***gt1 ra3* pre 3** | 94.65 | 24029388 | 21620296 | 90.0 | 18803100 | 87.0 |
| ***gt1 ra3* pre 4** | 94.75 | 28439898 | 25672623 | 90.3 | 22436350 | 87.4 |
| **A619 mid 1** | 94.88 | 28799650 | 25898540 | 89.9 | 22698302 | 87.6 |
| **A619 mid 2** | 94.95 | 25088136 | 22579052 | 90.0 | 19812404 | 87.7 |
| **A619 mid 3** | 94.62 | 26505978 | 23955111 | 90.4 | 21206660 | 88.5 |
| **A619 mid 4** | 94.91 | 24744422 | 22310002 | 90.2 | 19663897 | 88.1 |
| ***gt1* mid 1** | 95.13 | 26033539 | 23264791 | 89.4 | 20748143 | 89.2 |
| ***gt1* mid 2** | 95.39 | 27220133 | 24414794 | 89.7 | 22554173 | 92.4 |
| ***gt1* mid 3** | 94.86 | 32402821 | 29218207 | 90.2 | 26266340 | 89.9 |
| ***gt1* mid 4*** | 95.51 | 22241829 | 19709207 | 88.6 | 17373649 | 88.1 |
| ***ra3* mid 1** | 95.4 | 25852429 | 22846278 | 88.4 | 20268341 | 88.7 |
| ***ra3* mid 2** | 95.31 | 27493972 | 24510549 | 89.1 | 21807613 | 89.0 |
| ***ra3* mid 3** | 95 | 27835165 | 24828535 | 89.2 | 22159301 | 89.2 |
| ***ra3* mid 4** | 95.28 | 27337867 | 24651279 | 90.2 | 22090406 | 89.6 |
| ***gt1 ra3* mid 1** | 95.43 | 28575694 | 25480987 | 89.2 | 22526871 | 88.4 |
| ***gt1 ra3* mid 2** | 95.2 | 30921073 | 27566043 | 89.1 | 24604415 | 89.3 |
| ***gt1 ra3* mid 3** | 95.1 | 27722584 | 24813447 | 89.5 | 22140223 | 89.2 |
| ***gt1 ra3* mid 4** | 95.39 | 28295545 | 25333948 | 89.5 | 22602690 | 89.2 |

*not included in final analysis
